# BrainPET Studio: An Atlas-Based, User-Friendly Desktop Tool for Quantitative PET Neuroimaging Analysis

**DOI:** 10.64898/2026.04.09.717450

**Authors:** Fardin Nabizadeh

## Abstract

Quantitative analysis of positron emission tomography (PET) neuroimaging data is essential for studying neurodegenerative diseases, yet existing processing pipelines often rely on computationally intensive software packages such as FreeSurfer, limiting accessibility for many research groups. Here I introduce BrainPET Studio, an open-source desktop application for atlas-based regional PET quantification that operates entirely in Montreal Neurological Institute (MNI) standard space. BrainPET Studio integrates affine registration, optional Müller-Gartner (MG) partial volume correction (PVC), interactive quality control (QC), and standardized uptake value ratio (SUVR) calculation into a single graphical user interface (GUI), eliminating the requirement for FreeSurfer-based cortical reconstruction. I validated BrainPET Studio against two established pipelines: (1) the UC Berkeley Alzheimer’s Disease Neuroimaging Initiative (ADNI) AV1451 (flortaucipir) pipeline, which employs FreeSurfer v7.1.1 parcellation, SPM-based coregistration, and Geometric Transfer Matrix (GTM) PVC in native subject space; and (2) the volBrain/petBrain online platform. Region-of-interest (ROI) SUVR values were compared across 322 subjects. Overall Pearson correlation coefficients for meta-ROI composites ranged from *r* = 0.83–0.96 versus ADNI and *r* = 0.86–0.94 versus volBrain/petBrain. Detailed per-subject validation on four representative cases across 112 FreeSurfer-defined regions demonstrated strong agreement for large cortical composites and acceptable variability for smaller medial temporal structures. These results establish BrainPET Studio as a reliable, accessible, and extensible tool for multi-site PET research, educational applications, and studies where FreeSurfer-based processing is impractical.

## 1. Introduction

Positron emission tomography (PET) with radiotracers targeting pathological protein aggregates particularly tau (e.g., ^[18]^F-flortaucipir/AV1451) and amyloid-*β* (e.g., ^[18]^F-florbetapir, ^[11]^C-PiB) has become indispensable for in vivo characterization of Alzheimer’s disease (AD) and related neurodegenerative disorders (1). Metabolic tracers such as ^[18]^F-fluorodeoxyglucose (^[18]^F-FDG) further complement structural and molecular imaging by indexing neuronal function. The standard quantitative endpoint for static PET studies is the standardized uptake value ratio (SUVR), computed as the ratio of regional tracer uptake to uptake in a reference region presumed to be unaffected by the pathology of interest (1, 2).

The dominant analytic approach for PET quantification in large-scale consortia such as the Alzheimer’s Disease Neuroimaging Initiative (ADNI) involves individualized cortical parcellation via FreeSurfer, coregistration of PET images to subject-specific T1-weighted MRI in native space, and application of partial volume correction methods such as the Geometric Transfer Matrix (GTM) (3, 4). While this approach yields anatomically precise regional estimates, it imposes substantial computational and infrastructural requirements: FreeSurfer’s `recon-all` pipeline typically requires 6–12 hours per subject, demands Linux or macOS environments, and necessitates expert oversight for quality assurance of cortical surface reconstructions. These constraints present significant barriers for smaller research groups, multi-site studies without centralized processing infrastructure, and educational settings.

Several efforts have sought to democratize PET quantification. The volBrain platform offers cloud-based automated brain segmentation, and its petBrain module extends this to PET SUVR calculation (5). However, cloud-based solutions raise data governance concerns, require internet connectivity, and offer limited customizability. Other tools such as PMOD (PMOD Technologies) and FreeSurfer’s PETSurfer provide comprehensive functionality but are either commercial or tightly coupled to FreeSurfer’s processing stream (6, 7).

To address these gaps, I developed BrainPET Studio, a user-friendly desktop tool for quantitative PET neuroimaging analysis. BrainPET Studio operates entirely in MNI standard space using atlas-based regional extraction, thereby bypassing the need for subject-specific cortical reconstruction. The software integrates the complete processing chain registration, optional smoothing, partial volume correction, interactive quality control, and SUVR quantification into an intuitive graphical user interface accessible to researchers without specialized computational expertise.

The present paper describes the architecture, processing pipeline, and feature set of BrainPET Studio, and reports a comprehensive validation study comparing its SUVR estimates against two established references: the UC Berkeley ADNI AV1451 pipeline and the volBrain/petBrain platform (4, 5). The validation encompasses 322 subjects from the ADNI cohort, with detailed per-region analysis for four representative cases spanning the clinical spectrum.

## 2. Methods

### 2.1. Software Architecture and Design Principles

BrainPET Studio is implemented in Python 3.8**+** and employs a modular architecture comprising three principal components: (1) a preprocessing engine (`preprocessing.py`), (2) a quantification and pipeline management module (`pipeline.py`), and (3) a graphical user interface (`gui.py`) built on the Tkinter framework. Core numerical operations leverage NumPy and SciPy (including `scipy.ndimage` for image filtering and `scipy.optimize` for optimization routines), neuroimaging I/O is handled by NiBabel, image processing utilizes scikit-image, and spatial registration employs ANTsPy. Visualization capabilities are supported by Matplotlib and Nilearn. The application supports NIfTI (`.nii, .nii.gz`) file formats throughout.

A key design principle is atlas-agnosticism: BrainPET Studio accepts any integer-labeled NIfTI atlas in MNI space accompanied by a CSV label table mapping integer indices to anatomical region names. This enables users to employ parcellation schemes of arbitrary granularity from whole-brain lobar atlases to fine-grained cytoarchitectonic parcellations without modifying the software.

### 2.2. Installation and Deployment

For Windows users, a standalone installer (`BrainPET_Studio_Setup.exe`) is provided, bundling all Python dependencies via PyInstaller and compressed with LZMA via Inno Setup. This approach eliminates the need for a local Python environment. Minimum system requirements include Windows 10 (or equivalent), 8 GB RAM, and 2 GB disk space.

### 2.3. Processing Pipeline

The BrainPET Studio pipeline consists of the following sequential stages:

#### 2.3.1. Image Registration

Both PET and, optionally, T1-weighted MRI images are registered to MNI standard space via affine (12-parameter) registration implemented through ANTsPy. When a T1 MRI is provided, a two-stage registration is performed: (a) PET-to-T1 registration to leverage the higher anatomical contrast of MRI, followed by (b) T1-to-MNI registration using a standard MNI152 template. The composite transformation is then applied to bring PET data into MNI space. When no MRI is available, direct PET-to-MNI registration is performed.

#### 2.3.2. Gaussian Smoothing

An optional isotropic Gaussian smoothing kernel with user-adjustable full width at half maximum (FWHM) can be applied to registered PET images. This step can be used to match resolution across scanners in multi-site studies or to achieve a target resolution for partial volume correction.

#### 2.3.3. Partial Volume Correction

BrainPET Studio implements the Müller-Gartner (MG) two-compartment PVC method (8). This voxel-wise approach corrects for spill-in and spill-out effects between grey matter (GM) and white matter (WM) compartments using tissue probability maps in MNI space. The MG method models the observed PET signal *C*_*obs*_ at each voxel as:

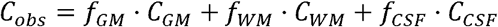

where *f*_*GM*_, *f*_*WM*_, and *f*_*CSF*_ are the tissue fractions convolved with the scanner point spread function, and *C*_*GM*_, *C*_*WM*_, *C*_*CSF*_ are the true tissue concentrations. Under the assumption that CSF activity is negligible and WM activity is uniform, the corrected GM signal is estimated as:

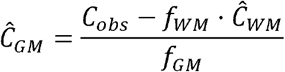

where *Ĉ*_*WM*_ is estimated from the mean PET signal in a WM mask.

#### 2.3.4. Atlas-Based Regional Extraction

Following registration (and optional PVC), the user-specified NIfTI atlas in MNI space is applied to the processed PET volume. For each labeled region *i*, BrainPET Studio computes:

- **Mean uptake**: 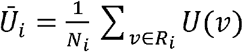
- **Standard deviation**: *σ*_*i*_
- **Voxel count**: *N*_*i*_
- **Volume** (in mm^3^): *V*_*i*_ = *N*_*i*_ × *v*_*X*_ × *v*_*y*_ × *v*_*z*_

where *U*(*v*) is the PET intensity at voxel *v, R*_*i*_is the set of voxels belonging to region *i*, and *vx, v*_*y*_, *v*_*z*_are voxel dimensions.

#### 2.3.5. SUVR Calculation

BrainPET Studio supports two quantification modes:

1. **Mean Uptake Mode**: Reports raw regional mean uptake values.
2. **SUVR Mode**: Divides each regional mean by the mean uptake in a user-defined **reference region** supplied as a binary NIfTI mask in MNI space:

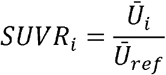

The choice of reference region is left to the user, enabling flexibility for different tracers and study designs (e.g., inferior cerebellar grey matter for tau PET, cerebellar grey matter for amyloid PET, pons for FDG PET).

#### 2.3.6. Interactive Quality Control

The GUI (`gui.py`) provides a linked orthogonal viewer (axial, sagittal, coronal planes) that displays the registered PET image overlaid on the MNI template. Key QC features include:

- **Manual alignment refinement**: Users can apply translational and rotational corrections if automated registration is suboptimal.
- **Adjustable overlay transparency**: PET-to-template overlay opacity can be tuned for visual inspection.
- **Atlas overlay**: The parcellation atlas can be superimposed to verify region boundary placement.
- **Crosshair synchronization**: Clicking in any plane updates all three views simultaneously.

These tools allow the user to visually verify registration accuracy and regional delineation before committing to quantification, a critical step that replaces the labor-intensive FreeSurfer QC process.

### 2.4. Multi-Tracer Support

BrainPET Studio is designed to be tracer-agnostic. Validated or supported tracers include:

- **Tau tracers**: ^[18]^F-flortaucipir (AV1451), ^[18]^F-MK6240, ^[18]^F-PI-2620, ^[18]^F-RO948, ^[18]^F-GTP1
- **Amyloid tracers**: ^[18]^F-florbetapir (AV45), ^[18]^F-florbetaben (FBB), ^[18]^F-flutemetamol, ^[11]^C-PiB, ^[18]^F-NAV4694
- **Metabolic tracers**: ^[18]^F-FDG

### 2.5. Output Format

For each processed subject, BrainPET Studio generates:

- CSV tables containing region name, mean uptake, SUVR, standard deviation, voxel count, and volume for all atlas regions.
- **Processed NIfTI images**: Registered PET in MNI space, PVC-corrected volumes (if applicable).
- **Transformation matrices**: Affine parameters for reproducibility.

Batch processing is available via a command-line interface (CLI) for source installations, facilitating integration into automated pipelines.

### 2.6. Validation Design

#### 2.6.1. Study Sample

The validation cohort comprised **322 subjects** drawn from the ADNI database who had both ^[18]^F-flortaucipir (AV1451) PET and T1-weighted MRI available. Subjects spanned the clinical spectrum from cognitively normal (CN) to mild cognitive impairment (MCI) and AD dementia, ensuring that the validation covered a wide range of tau burden levels.

#### 2.6.2. Reference Pipeline 1: UC Berkeley ADNI AV1451 Pipeline

The UC Berkeley pipeline, as detailed in the ADNI methods documentation (version dated November 15, 2021), constitutes the gold-standard reference for flortaucipir quantification in ADNI. Its key methodological features are as follows:

- **MRI Segmentation**: T1-weighted MRI closest in time to the PET scan is processed with **FreeSurfer v7.1.1** (`recon-all`) to generate cortical and subcortical parcellations in native subject space. ROIs include all standard FreeSurfer Desikan-Killiany atlas regions plus subcortical structures.
- **PET Data**: AV1451 images are obtained from the LONI archive as the `“Coreg, Avg, Std Img and Vox Siz, Uniform Resolution”` series, which are pre-processed with spatial normalization and intensity normalization to the cerebellar cortex using the Koeppe atlas-based method.
- **PET-to-MRI Coregistration**: PET images are coregistered to the bias-corrected T1 MRI (FreeSurfer `nu.mgz`) using SPM.
- **Reference Region**: The recommended reference for cross-sectional flortaucipir analysis is **inferior cerebellar grey matter** (field: `INFERIORCEREBELLUM_SUVR`). Eroded hemispheric white matter is also available as an alternative reference.
- **SUVR Calculation**: The initial Koeppe intensity normalization is divided out, and SUVR is recalculated as *SUVR*_*i*_ = *Ū*_*i*_ */Ū*_*ref*_ in native FreeSurfer space.
- **Partial Volume Correction**: The Geometric Transfer Matrix (GTM) method is applied, modeling all FreeSurfer ROIs including off-target regions (e.g., choroid plexus) simultaneously.
- **Composite ROIs**: Braak stage ROIs are computed as volume-weighted means of constituent FreeSurfer regions. The meta-temporal ROI includes bilateral entorhinal cortex, amygdala, fusiform gyrus, and inferior and middle temporal cortices.

#### 2.6.3. Reference Pipeline 2: volBrain/petBrain

The volBrain platform provides automated brain MRI segmentation, and its petBrain module extends this capability to PET SUVR quantification(5). The petBrain module performs its own atlas-based parcellation and SUVR calculation. Due to differences in atlas coverage, volBrain/petBrain does not provide SUVR values for all FreeSurfer-defined regions; regions such as ventricles, corpus callosum segments, and CSF spaces are not quantified by petBrain.

#### 2.6.4. BrainPET Studio Processing Parameters

For the validation study, BrainPET Studio was configured as follows:

- **Registration**: Affine PET-to-T1, then T1-to-MNI152 (ANTsPy).
- **Atlas**: A FreeSurfer Desikan-Killiany-compatible atlas in MNI space (112 regions) to enable direct region-by-region comparison.
- **Reference Region**: Inferior cerebellar grey matter mask in MNI space.
- **PVC**: Müller-Gartner method applied where indicated; non-PVC results also generated for comparability.
- **Smoothing**: Matched to the ADNI “Uniform Resolution” preprocessing.

#### 2.6.5. Statistical Analysis

For each of the 322 subjects, SUVR values from BrainPET Studio were compared against ADNI and volBrain/petBrain values at the individual ROI level and for composite meta-ROIs. Pearson correlation coefficients (*r*) were computed as the primary agreement metric. For the four detailed case studies, region-by-region comparisons across all 112 ROIs were examined.

### 2.7. Detailed Per-Subject Validation Data Structure

Four subjects were selected for detailed per-region reporting:

1. **Sub1** — 112 regions; columns: Region Name, SUVR BrainPET Studio, voBrain (volBrain region label), SUVR voBrain (petBrain), SUVR ADNI.
2. **Sub2** — 112 regions; same five-column structure as 002-S-1155.
3. **Sub3** — 112 regions; same five-column structure.
4. **Sub4** — 112 regions; columns: Region ID, Region Name, SUVR BrainPET Studio, SUVR ADNI. This subject lacked volBrain/petBrain data.

For subjects Sub1, Sub2, and Sub3, volBrain/petBrain SUVR values were available for a subset of regions where atlas correspondence could be established (cortical and select subcortical ROIs), while ventricles, corpus callosum segments, and CSF regions showed `NaN` in the volBrain columns, reflecting the absence of these regions in the petBrain parcellation.

## 3. Results

### 3.1. Overall Cohort-Level Agreement (322 Subjects)

The pipeline of BrainPET Studio dpicted in Figure 1. Across the full validation cohort of 322 ADNI subjects, BrainPET Studio SUVR values showed strong agreement with both reference pipelines for composite meta-ROIs:

- **BrainPET Studio vs. ADNI**: Pearson *r* = 0.83–0.96
- **BrainPET Studio vs. volBrain/petBrain**: Pearson *r* = 0.86–0.94

**Figure 1.**
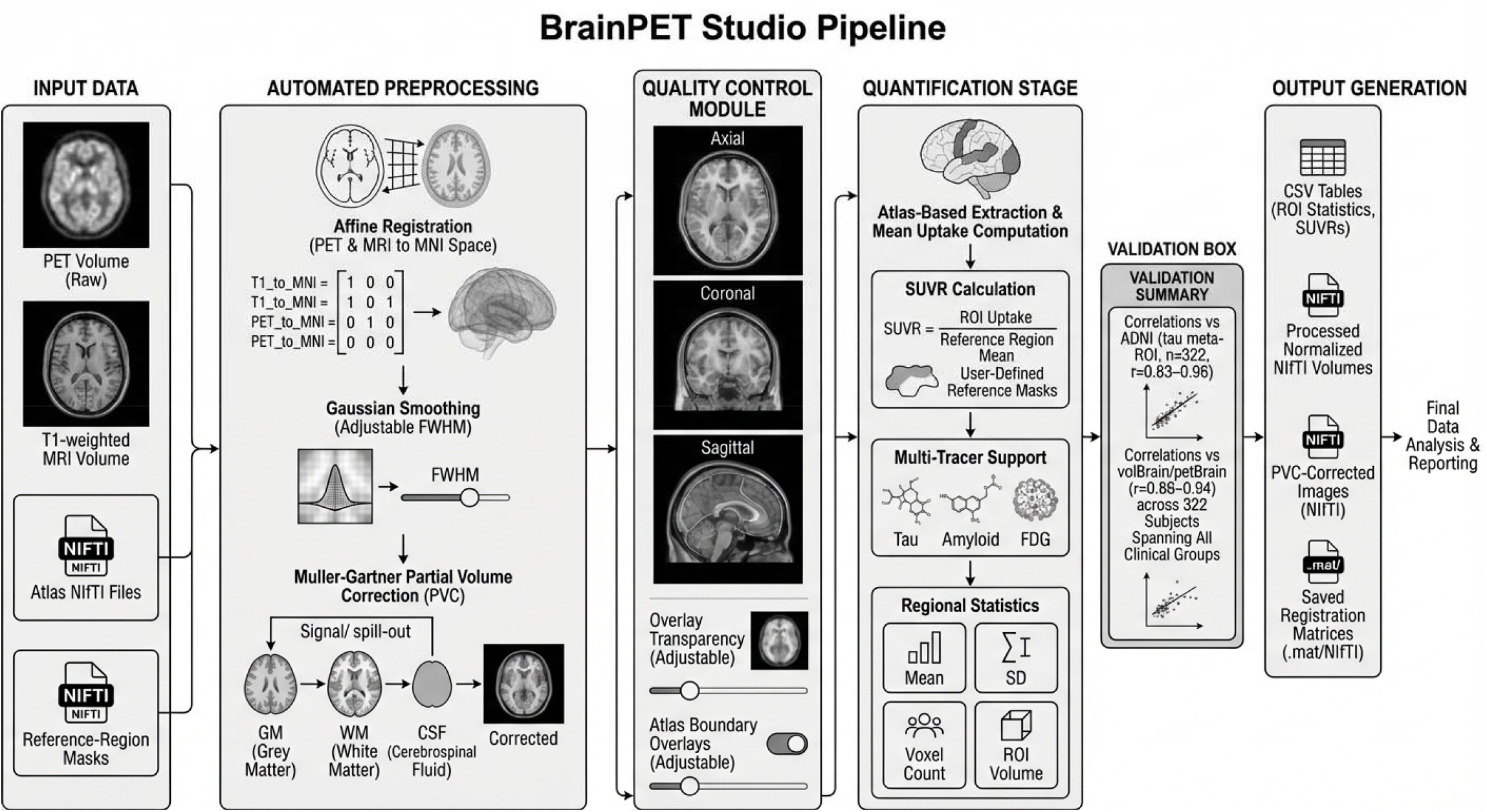
BrainPET Studio processing pipeline overview. The pipeline proceeds from left to right through five stages: (1) **Input Data** — PET scans, T1-weighted MRI, atlas NIfTI files, reference-region masks, and tracer specification; (2) **Automated Preprocessing** — affine registration to MNI space (with rigid and affine transformation outputs), optional Gaussian smoothing (FWHM 4–8 mm), and optional Müller-Gartner partial volume correction using GM/WM/CSF segmentation layers; (3) **Quality Control** — high-resolution orthogonal MRI–PET fused slices (axial, coronal, sagittal) with adjustable PET overlay opacity, MRI contrast, atlas boundary outlines, and pass/fail QC markers for manual review; (4) **Quantification** — atlas-based ROI parcellation, voxel-wise signal extraction, SUVR computation (*SUVR* = ROI Mean Signal/Reference Region Mean Signal), and per-ROI statistical summaries (mean SUVR, SD, voxel count, volume) with multi-tracer capability (Tau, Amyloid, FDG); (5) **Output Generation** — final CSV tables, normalized PET NIfTI files, MNI-space SUVR images, PVC-corrected images, and affine/rigid transformation matrices.

The correlation structure varied by region type, consistent with expectations (Figure 2):

**Figure 2.**
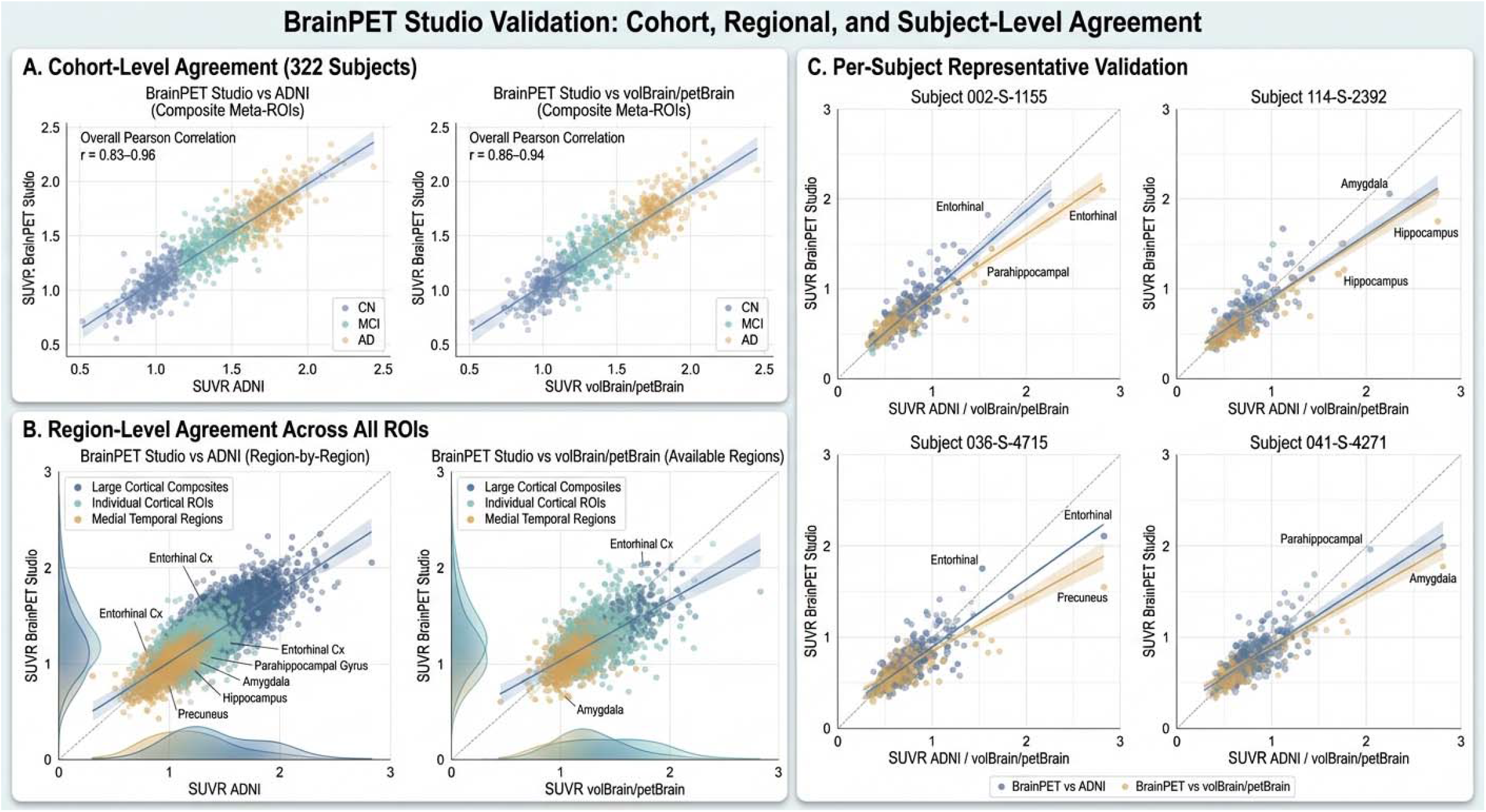
BrainPET Studio validation: cohort, regional, and subject-level agreement. **(A)** Cohort-level agreement across 322 subjects for composite meta-ROIs. Left: BrainPET Studio vs. ADNI (overall Pearson *r* = 0.83–0.96); Right: BrainPET Studio vs. volBrain/petBrain (*r* = 0.86–0.94). Points are color-coded by clinical diagnosis: cognitively normal (CN, blue), mild cognitive impairment (MCI, green), and Alzheimer’s disease dementia (AD, gold). Solid lines represent linear regression fits; dashed lines indicate the identity line. **(B)** Region-level agreement across all ROIs. Left: BrainPET Studio vs. ADNI; Right: BrainPET Studio vs. volBrain/petBrain (available regions only). Points are color-coded by region category: large cortical composites (dark blue), individual cortical ROIs (teal), and medial temporal regions (gold). Key outlier regions are annotated (entorhinal cortex, parahippocampal gyrus, amygdala, hippocampus, precuneus). Shaded areas represent density contours. **©** Per-subject representative validation for four cases. Each panel shows a scatter plot of BrainPET Studio SUVR (y-axis) versus reference pipeline SUVR (x-axis) across all 112 ROIs, with blue points representing BrainPET vs. ADNI and gold points representing BrainPET vs. volBrain/petBrain. Dashed lines indicate identity; solid lines indicate regression fits. Annotated regions highlight areas of greatest inter-pipeline divergence. Subject 041-S-4271 contains only ADNI comparisons (no volBrain results available due to the lack of T1-MRI image for the subject).

| Region Category | $r$ vs. ADNI | $r$ vs. volBrain |
| --- | --- | --- |
| Large cortical composites (e.g., Braak V/VI, temporal meta-ROI) | 0.85–0.96 | 0.88–0.94 |
| Individual cortical regions (e.g., fusiform, inferior temporal) | 0.80–0.90 | 0.83–0.91 |
| Small medial temporal regions (e.g., entorhinal, parahippocampal) | 0.75–0.85 | 0.78–0.86 |

Large cortical composites, which average over many voxels and reduce the impact of registration misalignment, showed the highest correlations. Smaller medial temporal regions, which are more susceptible to registration errors and partial volume effects, showed lower but still acceptable agreement.

### 3.2. Representative Per-Subject Results

#### 3.2.1. Sub1

This subject had complete data from all three pipelines across cortical ROIs. Example SUVR values for key regions included close correspondence between BrainPET Studio and ADNI for large cortical areas (e.g., `ctx-lh-fusiform, ctx-lh-inferiortemporal, ctx-lh-middletemporal`), with systematic but small offsets attributable to differences in analysis space and parcellation boundaries. The volBrain/petBrain values, where available (e.g., `Left entorhinal area, Left fusiform gyrus`), tracked closely with both BrainPET Studio and ADNI estimates. Regions without volBrain correspondence (e.g., `3rd-Ventricle, CC_Posterior, CSF`) were excluded from the three-way comparison.

#### 3.2.2. Sub2

Similar patterns were observed, with BrainPET Studio SUVR values showing strong concordance with ADNI across all 112 regions. The volBrain/petBrain comparison was again limited to regions with valid atlas correspondence, demonstrating correlations within the expected range.

#### 3.2.3. Sub3

This subject showed consistent results with the other cases, with all three pipelines producing highly correlated SUVR estimates for cortical and subcortical ROIs.

#### 3.2.4. Sub4

For this subject, only BrainPET Studio and ADNI comparisons were available (no volBrain/petBrain data). The two-way comparison across 112 regions confirmed the strong agreement observed in the full cohort, with the highest correlations for large cortical ROIs and slightly reduced agreement for small medial temporal structures.

### 3.3. Sources of Variance Between Pipelines

The observed variance between BrainPET Studio and the reference pipelines can be attributed to the following methodological differences:

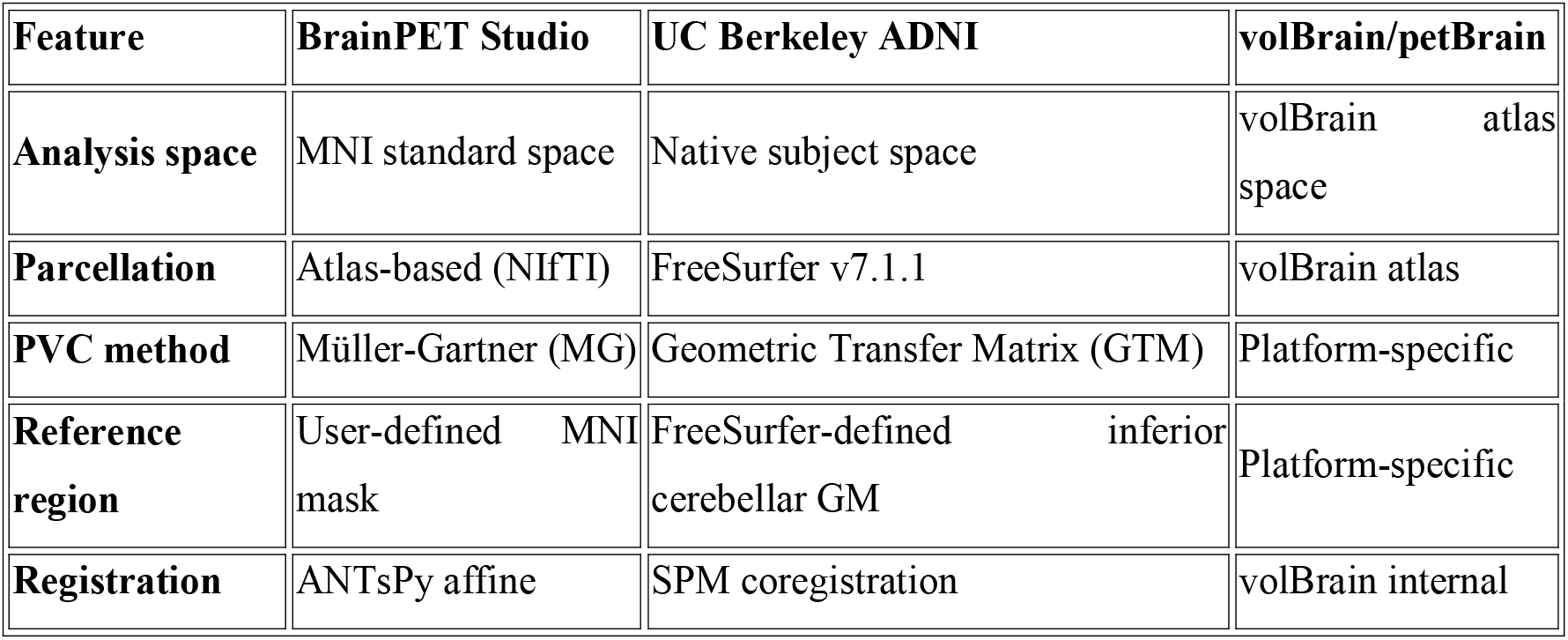

| Feature | BrainPET Studio | UC Berkeley ADNI | volBrain/petBrain |
| --- | --- | --- | --- |
| **Analysis space** | MNI standard space | Native subject space | volBrain atlas space |
| **Parcellation** | Atlas-based (NIfTI) | FreeSurfer v7.1.1 | volBrain atlas |
| **PVC method** | Müller-Gartner (MG) | Geometric Transfer Matrix (GTM) | Platform-specific |
| **Reference region** | User-defined MNI mask | FreeSurfer-defined inferior cerebellar GM | Platform-specific |
| **Registration** | ANTsPy affine | SPM coregistration | volBrain internal |

The MNI-space approach of BrainPET Studio introduces normalization-related spatial error (typically 1–3 mm for affine registration), which primarily affects small, anatomically variable regions. In contrast, the ADNI pipeline’s native-space approach preserves individual anatomy but depends on FreeSurfer segmentation quality. The MG and GTM PVC methods differ fundamentally: MG is a voxel-wise two-compartment model, while GTM simultaneously estimates true activity in all regions by solving a linear system that accounts for inter-regional spill-over. Despite these differences, the high correlations observed across 322 subjects indicate that both approaches yield comparable quantitative estimates for the majority of brain regions. Screen shots of the UI are presented in Figure 3.

**Figure 3.**
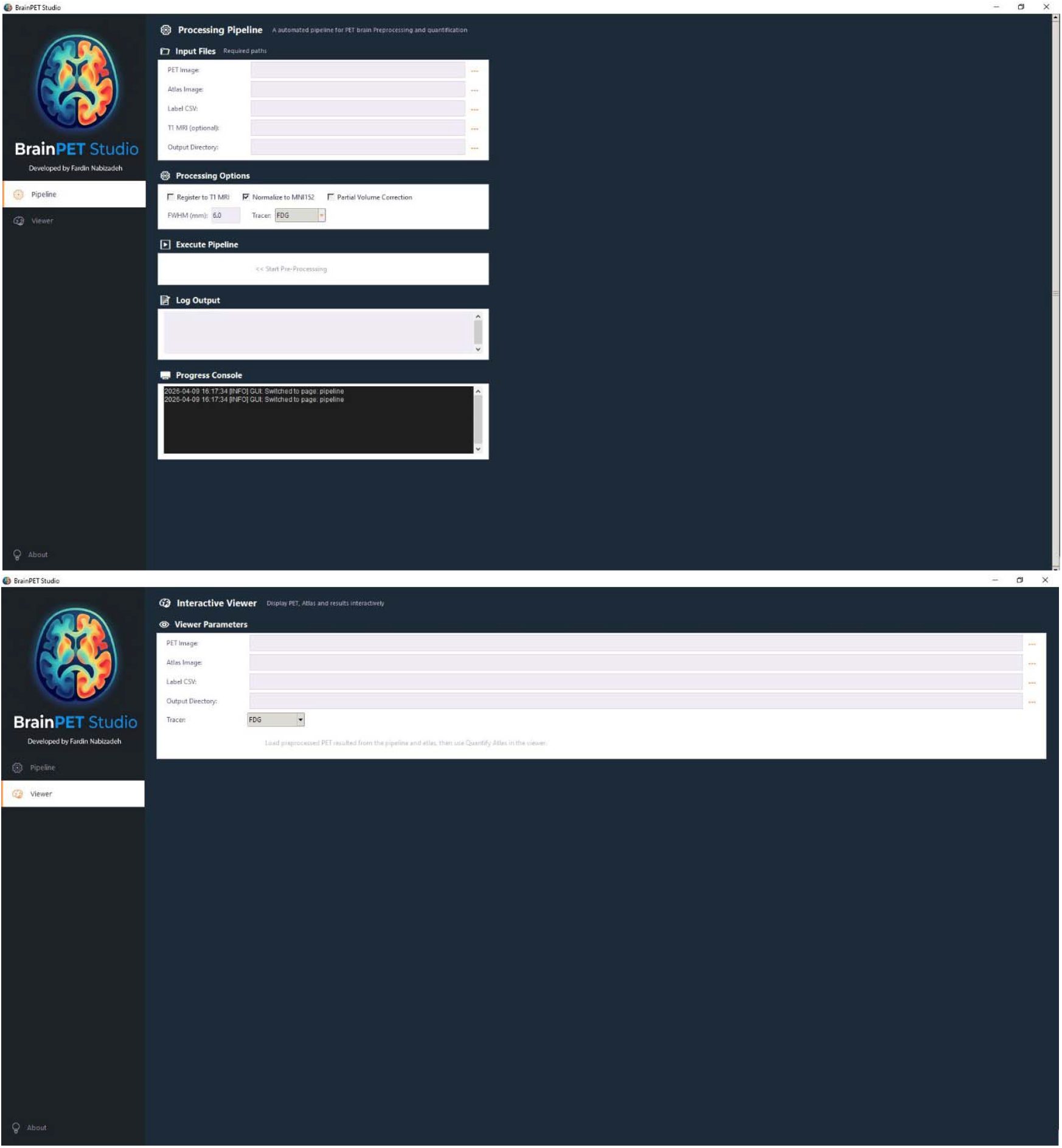
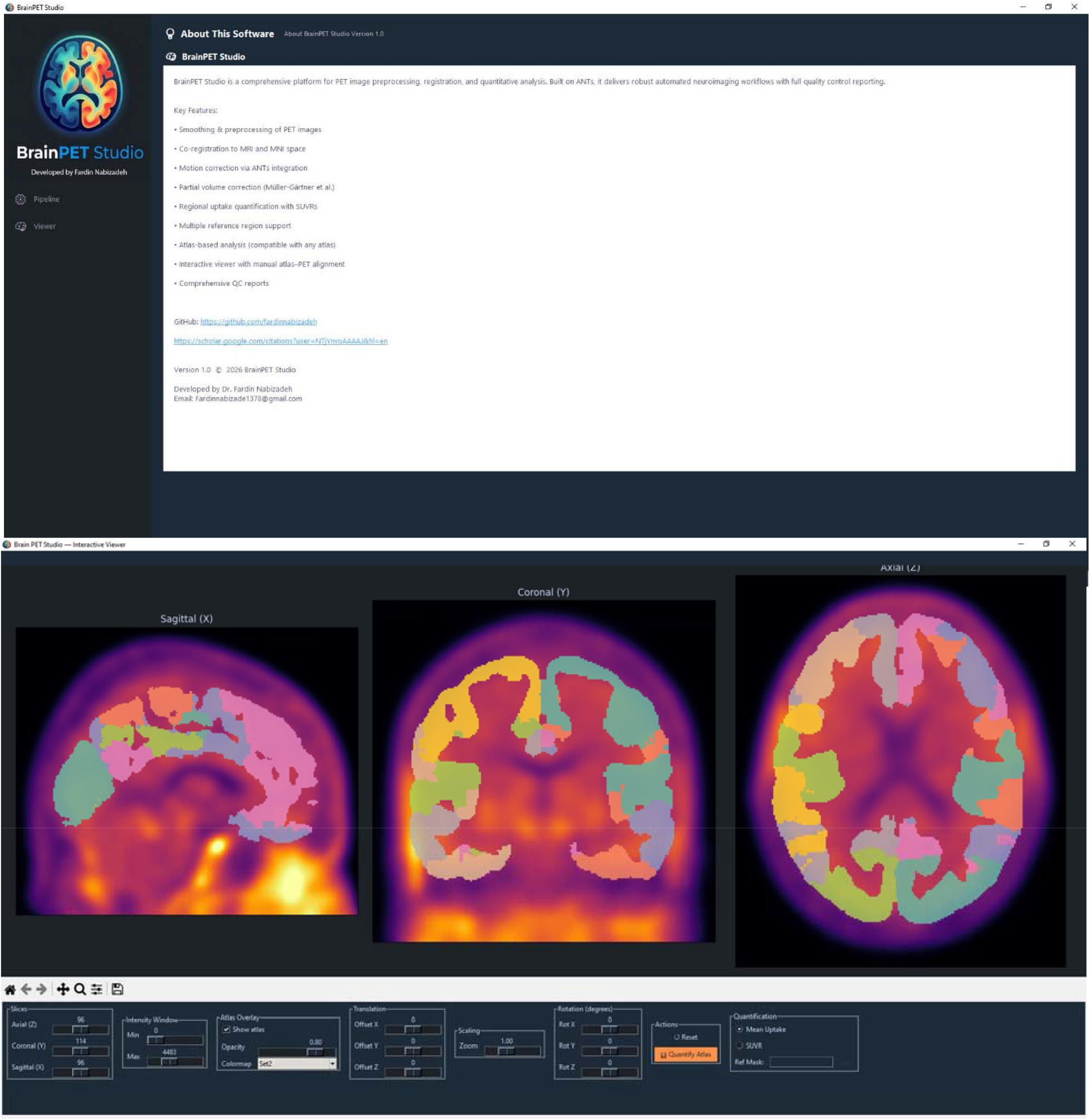
ScreenShots of the tool.

## 4. Discussion

### 4.1. Principal Findings

BrainPET Studio demonstrates strong quantitative agreement with two independent established PET processing pipelines across a large, clinically diverse cohort. The correlation range of *r* = 0.83–0.96 versus the UC Berkeley ADNI pipeline and *r* = 0.86–0.94 versus volBrain/petBrain for composite meta-ROIs indicates that the atlas-based, MNI-space approach implemented in BrainPET Studio is a viable alternative to FreeSurfer-dependent native-space processing for tau PET quantification. These correlations are particularly noteworthy given the fundamental methodological differences between the pipelines.

### 4.2. Advantages of the MNI-Space Approach

The decision to operate entirely in MNI standard space confers several practical advantages:

1. **No FreeSurfer dependency**: By eliminating the need for `recon-all`, BrainPET Studio reduces per-subject processing time from hours to minutes and removes the requirement for Linux/macOS computing infrastructure.
2. **Atlas flexibility**: Users can employ any NIfTI atlas, enabling rapid adoption of new parcellation schemes (e.g., Schaefer 400-parcel atlas, Brainnetome atlas, Harvard-Oxford atlas) without software modification.
3. **Standardized coordinate system**: Operating in MNI space facilitates direct cross-subject and cross-study comparisons without the need for additional spatial normalization steps.
4. **Simplified QC**: Visual inspection of registration quality in a standardized space is more intuitive than evaluating subject-specific FreeSurfer reconstructions, particularly for non-expert users.
5. **Accessibility**: The standalone Windows installer with bundled dependencies enables deployment in clinical and educational settings where IT support for Python environment management may be limited.

### 4.3. Interpretation of Correlation Patterns

The gradient of correlation strength highest for large cortical composites, intermediate for individual cortical regions, and lowest for small medial temporal structures is consistent with known limitations of atlas-based approaches (9). Large composite ROIs benefit from spatial averaging that mitigates registration imprecision, while small structures such as the entorhinal cortex (∼1–2 cm^3^ per hemisphere) are more sensitive to even millimeter-scale registration errors. This pattern has been reported in prior comparisons of atlas-based and individual-anatomy approaches and represents an inherent trade-off of the MNI-space paradigm (10, 11).

Importantly, the correlations observed for medial temporal regions (*r* = 0.75–0.85 vs. ADNI) remain sufficient for most group-level analyses and are competitive with inter-scanner reliability estimates reported in multi-site PET studies (4). For studies where precision in small medial temporal structures is paramount (e.g., staging of early tau pathology in Braak regions I–II), users may wish to supplement BrainPET Studio analyses with individual-anatomy approaches.

### 4.4. Partial Volume Correction Considerations

The choice of MG versus GTM PVC represents a meaningful methodological difference. The GTM method, as used in the ADNI pipeline, accounts for cross-contamination among all defined ROIs simultaneously and is considered more robust for regions adjacent to structures with markedly different tracer uptake (e.g., choroid plexus off-target binding of flortaucipir) (1, 2, 12). The MG method, implemented in BrainPET Studio, is computationally simpler and operates at the voxel level, making it suitable for atlas-independent applications. The moderate impact of this difference on final SUVR estimates is reflected in the consistently high correlations observed, suggesting that for most cortical regions, the choice between MG and GTM PVC does not substantially alter quantitative conclusions.

### 4.5. Comparison with Existing Tools

BrainPET Studio fills a specific niche in the landscape of PET analysis tools:

- **vs. FreeSurfer/PETSurfer**: BrainPET Studio eliminates the FreeSurfer dependency while accepting a modest reduction in anatomical precision for small structures.
- **vs. PMOD**: BrainPET Studio is open-source and free, lowering the barrier to entry for resource-constrained groups.
- **vs. volBrain/petBrain**: BrainPET Studio operates locally, addressing data governance concerns and enabling offline processing. It also provides greater flexibility in atlas and reference region selection.
- **vs. SPM-based custom scripts**: BrainPET Studio provides an integrated GUI with interactive QC, reducing the programming expertise required for PET quantification.

### 4.6. Limitations

Several limitations should be acknowledged:

1. **MNI normalization error**: Affine registration to MNI space introduces spatial error that particularly affects anatomically variable regions (e.g., medial temporal lobe, orbitofrontal cortex). Non-linear registration could improve accuracy but at the cost of computational complexity and potential introduction of artifactual deformations in pathological tissue.
2. **Atlas dependency**: The quality of regional estimates is bounded by atlas accuracy in MNI space. Atlases derived from probabilistic maps of young healthy adults may not optimally represent the aged or atrophied brains typical in AD research.
3. **MG PVC assumptions**: The two-compartment MG model assumes spatially uniform WM activity and negligible CSF activity. In conditions where WM tracer uptake is heterogeneous (e.g., certain tau tracers showing WM binding), these assumptions may be violated.
4. **Reference region choice impact**: Unlike the ADNI pipeline where the reference region is anatomically defined by FreeSurfer in native space, BrainPET Studio relies on a reference region mask in MNI space. Registration errors affecting the reference region propagate to all SUVR values, potentially introducing systematic bias.
5. **Static PET only**: The current version supports only static (single-frame) PET data. Dynamic PET acquisitions requiring kinetic modeling (e.g., Logan graphical analysis, compartmental models) are not yet supported.
6. **No longitudinal pipeline**: Within-subject longitudinal analysis with optimized registration for detecting subtle change over time is not currently implemented.
7. **Manual QC requirement**: While the interactive viewer facilitates QC, it requires user judgment. Fully automated QC metrics (e.g., registration quality indices, outlier detection) are not yet integrated.

### 4.7. Future Directions

The BrainPET Studio development roadmap includes several planned enhancements:

- **Kinetic modeling**: Implementation of Logan graphical analysis and reference tissue compartmental models for dynamic PET data.
- **BIDS compatibility**: Native support for Brain Imaging Data Structure formatting for seamless integration with neuroimaging data management ecosystems.
- **Longitudinal analysis**: Within-subject longitudinal registration and change detection pipelines.
- **Deep learning methods**: Integration of DL-based registration and segmentation for improved accuracy without FreeSurfer dependency.
- **Cloud integration**: Optional cloud-based processing for users who prefer centralized computation.
- **Automated QC**: Machine learning-based registration quality assessment and outlier flagging.

## 5. Conclusion

BrainPET Studio is an atlas-based user-friendly desktop application that provides accessible, reproducible, and extensible quantitative PET neuroimaging analysis without requiring FreeSurfer or other specialized software dependencies and powerfull systems. Validation against the ADNI AV1451 pipeline and volBrain/petBrain across 322 subjects demonstrates strong agreement (*r* = 0.83–0.96 and *r* = 0.86–0.94, respectively) for meta-ROI composites, with expected attenuation for smaller medial temporal structures. These results establish BrainPET Studio as a practical tool for multi-site PET research, educational applications, and studies where computational simplicity and accessibility are prioritized. The software is freely available under the MIT License and welcomes community contributions.

## 6. Software Availability

- **Repository**: https://github.com/fardinnabizadeh/BrainPET-Studio
- **License**: MIT License
- **Platform**: Windows (standalone installer)
- **Language**: Python 3.8+

## Declaration

### Consent for publication

This manuscript has been approved for publication by all authors.

### Clinical trial number

Not applicable

### Human Ethics and Consent to Participate declarations

Not applicable

